# The genomic landscape of polymorphic human nuclear mitochondrial insertions

**DOI:** 10.1101/008144

**Authors:** Gargi Dayama, Sarah B. Emery, Jeffrey M. Kidd, Ryan E. Mills

## Abstract

The transfer of mitochondrial genetic material into the nuclear genomes of eukaryotes is a well-established phenomenon. Many studies over the past decade have utilized reference genome sequences of numerous species to characterize the prevalence and contribution of nuclear mitochondrial insertions to human diseases. The recent advancement of high throughput sequencing technologies has enabled the interrogation of genomic variation at a much finer scale, and now allows for an exploration into the diversity of polymorphic nuclear mitochondrial insertions (NumtS) in human populations. We have developed an approach to discover and genotype previously undiscovered Numt insertions using whole genome, paired-end sequencing data. We have applied this method to almost a thousand individuals in twenty populations from the 1000 Genomes Project and other data sets and identified 138 novel sites of Numt insertions, extending our current knowledge of existing Numt locations in the human genome by almost 20%. Most of the newly identified NumtS were found in less than 1% of the samples we examined, suggesting that they occur infrequently in nature or have been rapidly removed by purifying selection. We find that recent Numt insertions are derived from throughout the mitochondrial genome, including the D-loop, and have integration biases consistent with previous studies on older, fixed NumtS in the reference genome. We have further determined the complete inserted sequence for a subset of these events to define their age and origin of insertion as well as their potential impact on studies of mitochondrial heteroplasmy.

## INTRODUCTION

The presence of mitochondrial DNA in the nuclear genomes of eukaryotes has been well established, and recent reports have shown that this transfer of genetic material is an ongoing evolutionary process (Mourier et al. 2001; Ricchetti et al. 2004; Hazkani-Covo and Covo 2008; Hazkani-Covo et al. 2010; Soto-Calderon et al. 2012). In humans, these nuclear insertions of mitochondrial origin (NumtS) have been estimated to occur at a rate of ∼5×10^−6^ per germ cell per generation (Leister 2005) and have been implicated directly in a number of genetic disorders (Willett-Brozick et al. 2001; Ahmed et al. 2002; Borensztajn et al. 2002; Turner et al. 2003; Goldin et al. 2004) while also indirectly hindering studies of mitochondrial diseases (Yao et al. 2008). A total of 755 NumtS have been identified in version hg19 of the human reference genome (Calabrese et al. 2012), although some portion of these have likely arisen through the duplication of previously inserted Numts. These fragments range in size from 39bps to almost the entire mitochondrial sequence and are thought to integrate themselves through a non-homologous end joining (NHEJ) mechanism during double-strand break repair (Blanchard and Schmidt 1996; Ricchetti et al. 1999). Over evolutionary time, many have been highly modified due to inversions, deletions, duplications, and displaced sequences, but some remain very well conserved relative to their parent mitochondria genome. While these fragments appear to be randomly selected from different regions of the mitochondria, an underrepresentation of the D-loop region has been reported, though why this is observed is currently unknown (Tsuji et al. 2012).

Several studies have previously looked at the enrichment of Numt insertions found in the human reference genome assembly relative to different genomic features. Some reports have suggested that Numt insertions tend to co-localize with repetitive elements (Mishmar et al. 2004; Tsuji et al. 2012), while others have found them to be under represented (Gherman et al. 2007). Some groups have further shown an under representation of repetitive elements nearby NumtS in humans but not flanking Numt insertions found in chimpanzees (Jensen-Seaman et al. 2009). In addition, there is evidence that numts preferably insert into open chromatin regions, typically near A+T oligomer sequences (Tsuji et al. 2012). As these studies are primarily based on older, fixed insertions in the human lineage, it is possible that they may have been confounded by evolutionary mutational processes that have occurred since the time of these insertions. As such, an investigation into more recent insertions is warranted in order to determine any insertion biases that may lead to a greater understanding of how this transfer of genetic material occurs.

Another important aspect of NumtS is their potential effect on studies of mitochondrial heteroplasmy, which are cell or tissue level differences in individual mitochondrial genomes due in part to the high rate of mutation within these sequences (Meyer et al. 2014). Low levels of heteroplasmy are typical in healthy individuals, and recent reports have determined that each person carries between 1 to 14 heteroplasmies (Cann et al. 2002; Hazkani-Covo and Covo 2008; Hajirasouliha et al. 2010; He et al. 2010; Tsuji et al. 2012; Ramos et al. 2013; Diroma et al. 2014; Ye et al. 2014). However, higher levels of heteroplasmy have been implicated in aging and various disease such as Leber’s hereditary optic neuropathy, diabetes, deafness, and even cancer (Wallace 1994; Avital et al. 2012; Gasparre et al. 2013; Ross et al. 2013). The presence of NumtS can confound the study and diagnosis of these diseases through the mistaken identification of nuclear-specific Numt mutations as heteroplasmy (Song et al. 2008; Yao et al. 2008). Computational and molecular approaches have been developed to help reduce the effect of NumtS on these studies (Goto et al. 2011; Ramos et al. 2013; Jayaprakash et al. 2014; Wolff 2014; Ye et al. 2014), but they only make use of known NumtS already present in the reference sequence and do not take into account recent insertions which may be still prevalent in a significant portion of the population.

In contrast to the many studies that have utilized NumtS present in the human reference sequence, there has been comparably little exploration into the landscape of polymorphic NumtS in human populations (Zischler et al. 1995; Thomas et al. 1996) and the largest such investigation to date has identified only 14 segregating events through investigation of the 1000 Genomes Project INDEL catalog (Lang et al. 2012). While rigorous, this analysis was limited in its ability to find novel insertion polymorphisms not present in the human genome reference due to the size of the sequence reads in which the variants could be discovered, resulting in the identification of only four such events. Here, we describe a new method, *dinumt*, for identifying numt insertions in whole genomes sequenced using paired-end sequencing technology, thus allowing for a greater sensitivity in identifying Numt variants of all sizes. We applied this method to 999 individuals from the 1000 Genomes (Genomes Project et al. 2012) and HGDP (Cann et al. 2002; Martin et al. 2014) projects and conducted an updated enrichment analysis using these polymorphic insertions. We further sequenced a subset of the polymorphic NumtS we discovered and examined them for their age, origin and sequence characteristics, and assessed their potential impact on ongoing studies of mitochondrial heteroplasmy.

## RESULTS

### Detecting numt insertions in whole genome sequences

Our discovery approach identifies clusters of read pairs mapping to both the nuclear and mitochondrial genomes and then examines these regions for potential insertion events (Fig. 1). This is similar in principle to earlier strategies designed to identify insertions of novel genetic material by finding clusters of one-end anchored reads (Kidd et al. 2008; Hajirasouliha et al. 2010) and to discovering mobile element insertions (Hormozdiari et al. 2010; Quinlan et al. 2010; Ewing and Kazazian 2011; Stewart et al. 2011; Keane et al. 2013), but requires that reads map to either the mitochondria or known reference NumtS to report a putative insertion (see methods). Nearby clusters are grouped together based on their orientation and distance from each other and the surrounding region is examined for split reads that map partially to both the chromosome and mitochondria, indicating the precise molecular breakpoint of the insertion. These sites can then be systematically genotyped across the entire sample set using a statistical framework similar to that previously developed for SNPs (Li 2011) to determine the insertion copy number in each individual.

**Figure 1:**
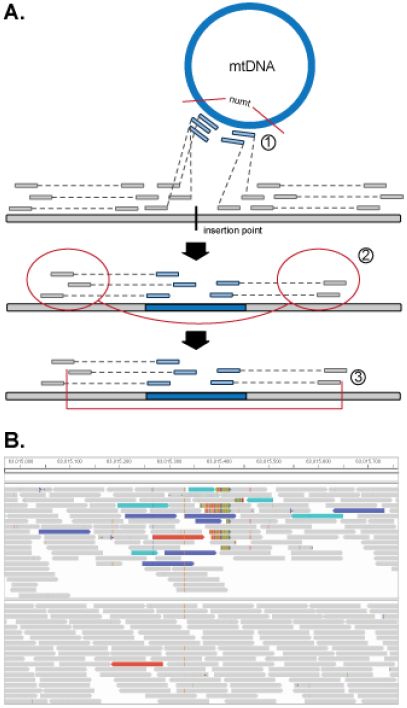
(A) Computational pipeline for Numt discovery: [1] Identification and filtration of paired reads with one read anchored to the mtDNA and another mapping to a nuclear chromosome; [2] Clustering and linking of nearby mapped nuclear reads together using insert size information; [3] Localization of insertion breakpoint using cluster distribution and truncated read alignments. (B) Example of Numt insertion (Poly_NumtS_2541) in sample HGDP00856 (top) compared to sample HGDP002222 (no insertion, bottom) on chromosome 8 as displayed in the IGV Browser. Sequences are represented by blocks and colored by the alignment of their mate sequence to the canonical location (grey), mitochondria genome (teal), or reference Numt homolog on chromosome 1 (blue). Multi-color bars represent split-reads whereby a portion of the sequence aligns to another location in the genome and are indicative of structural genomic breakpoints.

We applied our method to 946 low coverage, whole genomes that were sequenced in Phase 1 of the 1000 Genomes Project (Genomes Project et al. 2012) as well as 53 additional genomes sequenced to higher coverage from the HGDP (Cann et al. 2002; Martin et al. 2014) and were able to identify 141 polymorphic nuclear insertions of mitochondrial origin among all nuclear chromosomes except chrY, (Fig. 2), including 3 which had been previously characterized (Lang et al. 2012). No correlation was seen between the length of chromosome and the number of insertions and these insertions were fairly evenly distributed among all 20 different populations assessed (Supplemental Fig. S1, S2). On average, approximately 1.5 non-reference NumtS were seen in each sample and no significant bias was seen due to the sequence coverage (r^2^ = 0.21).

**Figure 2:**
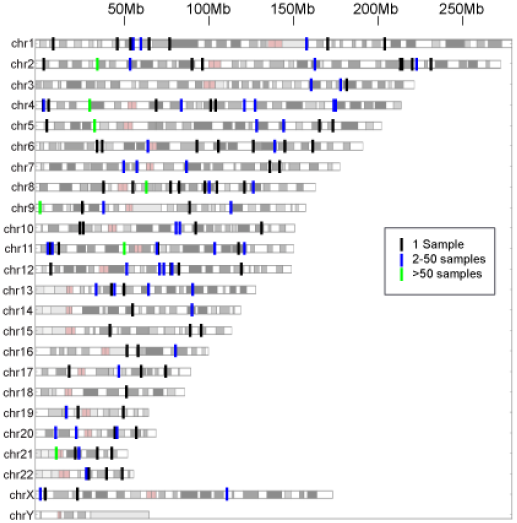
Chromosomal locations of individual Numt insertions, with color indicating its initial discovery in one sample (black), 2-50 samples (blue), and more than 50 samples (green).

We next assessed the overall accuracy of our approach. Using PCR, we were able to verify 23/24 of the predicted Numt insertion sites in the HGDP samples in which they were discovered and a further 17/18 from a subset of the lower coverage 1000 Genomes samples (Fig. 3A, Supplemental Table 1), with the events that we did not validate occurring either in segmental duplications or having uncertain breakpoints with potentially large insertion sizes, making them difficulty to conclusively verify. Additional validation with PCR panels across multiple samples showed a concordance for 713/748 (95.3%) of our predicted allele genotypes (Fig. 3B). We further validated that these were indeed of mitochondrial origin and not post-insertion duplications by Sanger sequencing through the breakpoints for 23 events (Supplemental Table 2. These results suggest that *dinumt* is able to accurately discover Numt insertions in whole genome sequence data.

**Figure 3:**
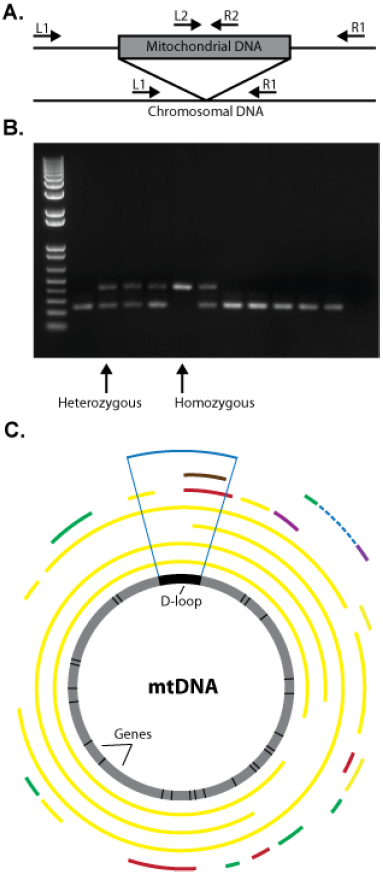
(A) PCR Strategy for validating Numt polymorphisms. Short insertions can be directly assessed using outer primers (L1, R1), while larger insertions require the additional use of internal primers (L1,R2; L2,R1). (B) Representative PCR validation panel for numt Poly_NumtS_1843 located on chromosome 4, with heterozygous and homozygous polymorphisms labeled with arrows. (C) Genomic context of sequenced polymorphic Numt insertions with respect to their Mitochondrial origin. Sequence identity of each Numt to the consensus mtDNA is indicated by color: 100% (green), 99% (yellow), 98% (purple), 97% (red), and 96% (brown). The D-loop region is denoted with vertical blue lines.

### Characteristics and Enrichment of Numt insertions

We conducted an analysis to confirm whether the insertion positions of these recent NumtS co-localized with specific genomic features, as has previously been assessed with the older, fixed events present in the reference genome (Supplemental Table 3). Using a series of permutation tests, we found no enrichment in regions containing CpG islands, microsatellites and other types of structural variants (P>0.05). Most Numt insertions were in intronic (42%) and intergenic regions (43%), consistent with expectations from random sampling, and we observed no insertions into coding exons. NumtS were also found in the 5’ and 3’ UTR’s as well as promoter and terminator regions (5Kbp up and downstream, respectively), albeit at a much lower frequency. Consistent with previous reports, we found a significant enrichment near repetitive regions (P<=0.004) (Tsuji et al. 2012) and an insertion preference for slightly higher %GC regions overall (Supplemental Fig. S3).

Interestingly, we neither observed enrichment for A+T oligomers immediately adjacent to the polymorphic insertions (Supplemental Fig. S4) nor a preference for open chromatin regions in the cell lines we investigated (P>0.05), as had been previously reported (Tsuji et al. 2012). This enrichment was prevalent even when limited to those NumtS for which we had validated breakpoints. To verify the consistency of our permutation approach, we applied the same analysis to the 610 reference NumtS described in Tsuji et al as well as the human-specific NumtS described in Lang et al (Lang et al. 2012) and were able to replicate their results. This difference at the insertion sites could represent a *bona fide* change in the integration mechanism for recent NumtS, but may also be an artifact from the way reference NumtS have been annotated relative to the mitochondria genome sequence.

### Analysis of Polymorphic Numt Sequences

While the precise insertion location for each Numt is informative, the underlying sequence itself can provide additional information. Using either direct Sanger sequencing of PCR products or subsequent primer walking and assembly for larger insertions (see methods, Supplemental Table 1), we were able to determine the sequence for 23 Numt insertions (Supplemental Table 2). Most NumtS sequences were small (<500bp), however we did identify a number of larger events including an almost complete mitochondrial genome insertion of 16,106bps in sample HGDP01275. We observed fragments originating from all parts of the mitochondrial genome, including multiple sequences overlapping the D-loop region (Fig. 2C) which had been previously reported as underrepresented in the human reference (Tsuji et al. 2012). Interestingly, these polymorphic sequences exhibited a higher %GC than that of their parent mitochondria genome (47% vs. 44%) and also showed similar characteristics to other fixed human-specific NumtS in the reference genome (Supplemental Fig. S5). In contrast, non-human specific NumtS in the reference showed a markedly lower %GC that is more consistent with the average nuclear genome %GC of 41.5%.

Using these sequences, we estimated when these insertions occurred in the human lineage by comparing the human mtDNA reference sequence to an inferred ancestral mitochondrial sequence and identifying diagnostic mutations that matched specific positions in each Numt sequence (see methods). We then used the fraction of alleles that matched those from the modern human mitochondrial sequence to derive an approximate age for each insertion, relative to an estimated human-chimpanzee divergence time of 6 million years (Table 1). We observed that most of the polymorphic insertions occurred within the past 1 million years, however there were six NumtS that were markedly older, including two that appear to have inserted over 2.5 million years in the past. We next constructed maximum-likelihood trees to compare fixed human-specific NumtS present in the reference genome (Fig. 4A) with the discovered polymorphic NumtS (Fig. 4B). As expected, the ongoing polymorphisms co-localized with the human lineage while the fixed events were likely inserted further back in time.

**Figure 4:**
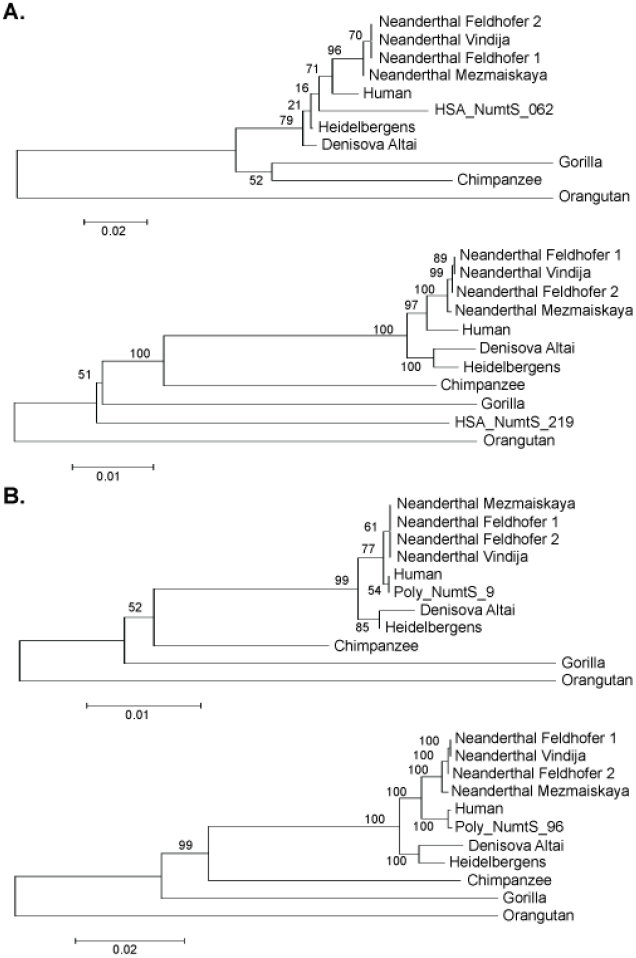
Phylogenetic trees for select (A) fixed and (B) polymorphic Numt insertion sequences relative to various species along the human and other primate lineages. Fixed NumtS were previously identified as human-specific (Jensen-Seaman et al. 2009) and are present in the human reference sequence (hg19). Polymorphic sequences were chosen from among the longest insertions that were identified. Bootstrap values are indicated at branch locations.

**Table 1.** Age estimations of sequenced Numt insertions using a consensus ancestral mitochondrial sequence

| Numt ID | Length of Insertion | # of Diagnostic Positions | # of Positions Matching Human | % of Human mtDNA | <sup>a</sup>Estimated Age of Insertion (MYA) |
| --- | --- | --- | --- | --- | --- |
| Poly_NumtS_139 | 391 | 20 | 20 | 100 | <0.1 |
| Poly_NumtS_1239 | 246 | 8 | 7 | 88 | 0.72 |
| Poly_NumtS_1259* | 547 | 40 | 37 | 92 | 0.48 |
| Poly_NumtS_1440 | 144 | 9 | 9 | 100 | <0.1 |
| Poly_NumtS_1843 | 163 | 11 | 10 | 91 | 0.54 |
| Poly_NumtS_1900 | 166 | 5 | 3 | 60 | 2.4 |
| Poly_NumtS_1929 | 229 | 8 | 7 | 88 | 0.72 |
| Poly_NumtS_2010 | 153/147<sup>c</sup> | 16 | 15 | 94 | 0.36 |
| Poly_NumtS_2186* | 245 | 0 | <sup>b</sup>NA | - | - |
| Poly_NumtS_2289* | 11840 | 577 | 426 | 86 | 0.84 |
| Poly_NumtS_2377 | 196 | 8 | 8 | 100 | <0.1 |
| Poly_NumtS_2541 | 1412 | 68 | 50 | 74 | 1.56 |
| Poly_NumtS_2578* | 16091 | 678 | 664 | 98 | 0.12 |
| Poly_NumtS_2611 | 119 | 6 | 6 | 100 | <0.1 |
| Poly_NumtS_2653 | 1665 | 91 | 69 | 76 | 1.44 |
| Poly_NumtS_316* | 655 | 2 | 2 | 100 | <0.1 |
| Poly_NumtS_430* | 482 | 39 | 29 | 74 | 1.56 |
| Poly_NumtS_445 | 145 | 3 | 3 | 100 | <0.1 |
| Poly_NumtS_531 | 269 | 17 | 16 | 94 | 0.36 |
| Poly_NumtS_709 | 140 | 7 | 3 | 43 | 3.42 |
| Poly_NumtS_1465* | 6775 | 261 | 256 | 98 | 0.12 |
| Poly_NumtS_1480* | 13783 | 580 | 447 | 77 | 1.38 |
| Poly_NumtS_1583 | 126 | 6 | 5 | 83 | 1.02 |

<sup>a</sup>Based on approximate divergence between human and chimpanzee of 6 million years (Meyer et al. 2014)
<sup>b</sup>Poly\_NumtS\_2186 did not overlap any of the identified diagnostic differences between the human and ancestral mitochondria
<sup>c</sup>Poly\_NumtS\_2010 had two noncontiguous fragments of lengths 153 and 147
<sup>\*</sup>Contains hypervariable D-loop region

### Assessment of Numt Impact on Heteroplasmy

Although many recent studies of mitochondrial heteroplasmy have taken NumtS into consideration (He et al. 2010; Li et al. 2010; Ramos et al. 2013; Bintz et al. 2014; Diroma et al. 2014; Hodgkinson et al. 2014; Jayaprakash et al. 2014), they have all been limited to those insertions present in the reference sequence. To assess the potential impact of more recent insertions, we compared our set of sequenced polymorphic NumtS to these studies by identifying single nucleotide differences in the Numt insertions relative to the mtDNA reference and comparing the allelic changes to those reported (Table 2). We identified 59 positions of possible Numt confounding, most of which occur in polymorphic insertions common in the general human population (MAF > 0.01). The samples used in most of these studies differ from those analyzed here, making direct inferences regarding these effects of NumtS difficult. However, one study (Diroma et al. 2014) had an intersecting set of individuals with our analysis, and we were able to determine that there were 8 positions within NumtS that were genotyped in those samples and had alleles matching the reported heteroplasmy.

**Table 2.** Examples of Numt insertion alleles matching identified mitochondrial heteroplasmic positions

| Position | mtDNA Allele | Numt Allele | Numt ID(s) | <sup>a</sup>Max Allele Frequency | <sup>b</sup>Study Ref. |
| --- | --- | --- | --- | --- | --- |
| 73 | A | G | Poly_NumtS_1465;Poly_NumtS_1480 | 0.026 | 1,5,7 |
| 263 | A | G | Poly_NumtS_1465;Poly_NumtS_1480 | 0.026 | 7 |
| 489 | T | C | Poly_NumtS_1465 | 0.007 | 5,7 |
| 750 | A | G | Poly_NumtS_1465;Poly_NumtS_1480 | 0.026 | 7 |
| 1438 | A | G | Poly_NumtS_1465;Poly_NumtS_1480 | 0.026 | 7 |
| 2706 | A | G | Poly_NumtS_1465;Poly_NumtS_1480 | 0.026 | 5,7 |
| 4769 | A | G | Poly_NumtS_2289 | 0.003 | 7 |
| 5460 | G | A | Poly_NumtS_2289 | 0.003 | 2,5 |
| 7028 | C | T | Poly_NumtS_2289 | 0.003 | 5,7 |
| 7220 | T | C | Poly_NumtS_2541 | 0.154 | 5 |
| 7256 | C | T | Poly_NumtS_2289 | 0.003 | 7 |
| 7521 | G | A | Poly_NumtS_2289;Poly_NumtS_2541 | 0.154 | 7 |
| 7861 | T | C | Poly_NumtS_2541 | 0.154 | 6,7 |
| 7912 | G | A | Poly_NumtS_2377 | 0.016 | 3 |
| 7927 | C | T | Poly_NumtS_2541 | 0.154 | 5 |
| 8021 | A | G | Poly_NumtS_2377 | 0.016 | 1 |
| 8122 | A | G | Poly_NumtS_2541 | 0.154 | 5 |
| 8152 | G | A | Poly_NumtS_2377 | 0.016 | 2 |
| 8206 | G | A | Poly_NumtS_2541 | 0.154 | 7 |
| 8251 | G | A | Poly_NumtS_2541 | 0.154 | 5 |
| 11176 | G | A | Poly_NumtS_1900 | 0.023 | 7* |
| 12612 | A | G | Poly_NumtS_2653 | 0.003 | 5,6,7 |
| 12630 | G | A | Poly_NumtS_2653 | 0.003 | 5 |
| 12705 | C | T | Poly_NumtS_2653;Poly_NumtS_1480 | 0.026 | 5,7 |
| 13506 | C | T | Poly_NumtS_2653 | 0.026 | 5,7 |
| 13650 | C | T | Poly_NumtS_2653 | 0.026 | 7 |
| 14766 | C | T | Poly_NumtS_1465;Poly_NumtS_1480 | 0.026 | 5,7 |
| 15043 | G | A | Poly_NumtS_1465 | 0.007 | 7 |
| 15301 | G | A | Poly_NumtS_1465 | 0.007 | 5,6,7 |
| 15326 | A | G | Poly_NumtS_1465;Poly_NumtS_1480 | 0.026 | 7 |
| 15575 | G | C | Poly_NumtS_1480 | 0.026 | 5 |
| 16093 | T | C | Poly_NumtS_1259 | 0.013 | 1,2,4,5,7* |
| 16129 | G | A | Poly_NumtS_430 | 0.568 | 4,7* |
| 16189 | T | A | Poly_NumtS_430 | 0.568 | 4 |
| 16209 | T | C | Poly_NumtS_1259 | 0.013 | 6,7 |
| 16218 | C | T | Poly_NumtS_430 | 0.568 | 4,7* |
| 16223 | C | T | Poly_NumtS_1259;Poly_NumtS_1465;Poly_NumtS_1480 | 0.026 | 2,4,5,6,7 |
| 16230 | A | G | Poly_NumtS_1259;Poly_NumtS_430 | 0.568 | 4,7 |
| 16234 | C | T | Poly_NumtS_1259 | 0.013 | 5 |
| 16249 | T | C | Poly_NumtS_430 | 0.568 | 4,7 |
| 16259 | C | A | Poly_NumtS_430 | 0.568 | 4 |
| 16263 | T | C | Poly_NumtS_430 | 0.568 | 4 |
| 16264 | C | T | Poly_NumtS_430 | 0.568 | 4 |
| 16274 | G | A | Poly_NumtS_430 | 0.568 | 4 |
| 16278 | C | T | Poly_NumtS_1259;Poly_NumtS_430 | 0.568 | 4,5,7* |
| 16284 | A | G | Poly_NumtS_430 | 0.568 | 4 |
| 16288 | T | C | Poly_NumtS_430 | 0.568 | 4 |
| 16290 | C | T | Poly_NumtS_430 | 0.568 | 4 |
| 16293 | A | C | Poly_NumtS_430 | 0.568 | 4,5 |
| 16301 | C | T | Poly_NumtS_430 | 0.568 | 4 |
| 16311 | T | C | Poly_NumtS_1259;Poly_NumtS_430 | 0.568 | 4,5,6,7* |
| 16355 | C | T | Poly_NumtS_1259;Poly_NumtS_430 | 0.568 | 4,5 |
| 16356 | T | C | Poly_NumtS_1259;Poly_NumtS_430 | 0.568 | 4 |
| 16362 | T | C | Poly_NumtS_1259 | 0.013 | 2,5,7 |
| 16368 | T | C | Poly_NumtS_430 | 0.568 | 4 |
| 16390 | G | A | Poly_NumtS_430 | 0.568 | 4,7* |
| 16519 | T | C | Poly_NumtS_1259;Poly_NumtS_430 | 0.568 | 5,7* |
| 16527 | C | T | Poly_NumtS_1259;Poly_NumtS_430 | 0.568 | 5 |

<sup>a</sup>Largest Numt allele frequency is listed from among sites with multiple overlapping insertions
<sup>b</sup>Studies included are 1:(He et al. 2010) , 2:(Li et al. 2010), 3:(Ramos et al. 2013), 4:(Bintz et al. 2014), 5:(Hodgkinson et al. 2014), 6:(Jayaprakash et al. 2014), and 7:(Diroma et al. 2014). It should be noted that (5) identified heteroplasmy in RNA sequences while the remaining studies were from DNA.
\*Matches sample specific heteroplasmy allele as reported in (Diroma et al. 2014)

## DISCUSSION

Almost every eukaryotic species that has had their genome fully sequenced to date has exhibited evidence for the transfer of organelle DNA to the nuclear genome. These NumtS occur in both animals and plants, and show a strong correlation with genome size and the total number of NumtS observed (Hazkani-Covo et al. 2010). In humans, there are approximately 755 annotated NumtS in the reference genome (Calabrese et al. 2012), though this number is variable depending on the methods and parameters used to identify their presence. However, very few of these are due to recent insertions and almost all NumtS that have been identified are present in every human genome. Indeed, only 14 events differentially present in human populations have been previously reported (Lang et al. 2012).

Here, we present a large-scale analysis of polymorphic mitochondrial insertions into the nuclear genome of humans. These recent Numt insertions share many characteristics with previously identified human-specific NumtS that are fixed in the genome, including their patterns of integration within the genome and sequence composition. We identified many NumtS that contain the mitochondrial D-loop, a noncoding region of the mitochondria that controls the synthesis of DNA and RNA within the organelle and typically exhibits a higher mutation rate than the rest of the genome. This region is often used in forensic and population genetics due to its two hypervariable regions that provide distinguishing polymorphisms between individuals (Budowle et al. 1999; Szibor et al. 2000) and has been previously found to be depleted among NumtS present in the human reference sequence (Tsuji et al. 2012). Previous studies have found little effect of existing, older NumtS on these types of assays (Goios et al. 2006), but have not taken into account these more recent insertions that are more likely to cause off-target amplification and erroneous conclusions. Our methods and data sets should thus provide a useful resource in these types of analyses.

The polymorphic nature of these insertions within the human lineage indicates that they have likely occurred since the most recent common ancestor between humans and chimpanzees, however it is unknown where they are relative to other species of humans such as Neanderthals or Denisovans. We thus attempted to date our insertions both through direct comparison with a consensus ancestral mitochondrial as well as through phylogenic analysis and found that most were integrated into the nuclear genome within the past 100,000 years. Over half of the NumtS we discovered were present in very low frequencies across the samples we interrogated (MAF <0.1%), suggesting that they were likely integrated even more recently than the resolution of our analysis would allow. This supports the theory that mitochondria gene transfer to the human nuclear genome is ongoing and prevalent (Ricchetti et al. 2004; Hazkani-Covo et al. 2010)

NumtS have been previously implicated in a number of sporadic disease cases through their integration into functional regions of the genome (Borensztajn et al. 2002; Turner et al. 2003; Goldin et al. 2004). While we did not identify any NumtS that would directly affect the coding region of a gene, we did identify a Numt insertion in a single individual that represented an almost entire insertion of a mitochondrial genome into chromosomal DNA (Fig. 5). This insertion was 16,106bps in size and integrated into a potential regulatory region in the first intron of the SDC2 gene (Kent et al. 2002), a member of the syndecan family that encodes an integral membrane protein and has been associated with cell proliferation and migration, including altered expression in several cancer cells (De Oliveira et al. 2012; Oh et al. 2013). We investigated whether this insertion may have had an effect on the expression of this gene by looking at recently published RNA-Seq data over these same samples (Martin et al. 2014), however this gene is not expressed in the tissues which were used in that analysis and so we are unable to draw any conclusions regarding its potential impact. It is tempting to speculate, however, that an insertion that is highly enriched for functional regions could indeed affect canonical gene structure and expression, and ongoing studies in individual tissues from projects such as GTEx (G. TEx Consortium 2013) and others may provide they keys for further investigating these types of events.

**Figure 5:**
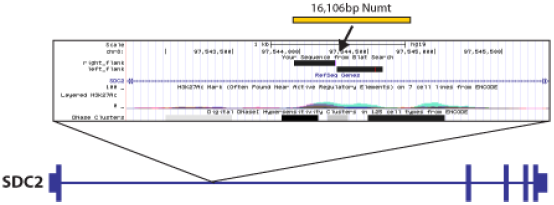
Insertion of an almost full-length mitochondrial insertion (Poly_NumtS_2578) into the first intron of the SDC2 gene in sample HGDP01275. Zoomed panel shows UCSC Genome Browser (http://genome.ucsc.edu) view of 3kbp surrounding region of insertion, with sequenced flanking regions of the insertion breakpoint indicated by solid black rectangles.

Finally, we explored the potential effect of our NumtS on studies of mitochondrial heteroplasmy and identified a number of positions within the mitochondrial genome that could be erroneously attributed to mutations in NumtS. It is possible that these heteroplasmies are prevalent and the allelic changes in the NumtS occurred prior to their insertion, and indeed a recent study using RNA-Seq expression data to examine heteroplasmy identified a number of these same positions (Hodgkinson et al. 2014). While it is possible that NumtS may be expressed at some low level in the nucleus, it is more likely that the reported heteroplasmies are bona fide mitochondrial differences. However we believe that our set of genotypes and insertion sequences will be a useful resource for future studies into mitochondrial heterogeneity.

## METHODS

### Data sources

Whole genome sequences were generated as a part of Phase I of the 1000 Genomes Project (http://www.1000genomes.org) with an average 4-6X sequence coverage and from the CEPH-Human Genome Diversity Project (HGDP,SRA: SRP036155) (Cann et al. 2002; Martin et al. 2014) with a higher average coverage of 5-20X. Alignments to version GRCh37/hg19 of the human reference genome were provided in BAM format and optimized using the Genome Analysis Toolkit (GATK) (McKenna A, 2010) and Picard (http://picard.sourceforge.net/), as described elsewhere (Genomes Project et al. 2012; Martin et al. 2014).

### Detection of Numt insertions

Non-reference NumtS were discovered in paired-end, whole genome sequences using a newly developed software package named *dinumt*, as outlined in Fig. 1. This approach first derives an empirical insert size distribution from the observed alignment positions of each read pair and then identifies sequences where one end aligns to either the mtDNA or a known reference Numt and the other maps elsewhere in the nuclear genome. These sequences are then clustered together based on their shared mapping orientation (forward or reverse) and whether or not they are within a distance of W_L_ from each other, where W_L_ is calculated as the derived mean_insert_length + 3 * insert_standard_deviation. Clusters are further linked together if they are within a distance of 2 * W_L_ from each other and are in the correct orientation relative to each other (forward to reverse). Individual sequence reads are then examined within the clusters to identify soft-clipped reads with breaks at the same position to identify putative breakpoint locations. The likelihood of an insertion is then calculated as

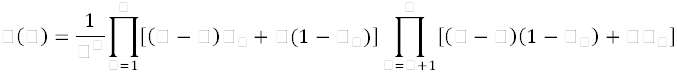

where *m* is the ploidy, *l* is the total number of reference supporting reads, *k* is the total number of insertion supporting reads, and *e* is the mapping error for read *j*, as modified from (Li 2011). Putative insertions were then filtered for quality (at least 50 using Phred scaled maximum insertion likelihood of non-reference allele), the number of total reads supporting the insertion (at least 4), and the depth of total coverage at the insertion point (at least 5).

Identified Numt insertions were then genotyped across the entire set of samples to identify sites which may have been previously missed or filtered in those individuals during the discovery step as well as to determine the copy number of the insertion. This was done by systematically examining each sample at an insertion location for clusters of reads supporting an insertion. In order to refine breakpoint positions, these clusters are used to search for positions where soft-clipped reads are consistently broken at the same location across samples and where the longest unaligned portion of such reads at that position map to the mtDNA reference genome. These refined positions are then utilized to determine how many sequences overlap the insertion location in an unbroken manner, supporting the reference allele, and how many contribute to the inserted Numt. These are then tabulated, combined with the read pair information, and scored in a similar fashion as the discovery step above. However, the reported genotype here reflects the overall maximum likelihood of that genotype (0/0, 0/1, 1/1) and not just that of an insertion (as above). Global genotypes are then used to construct population level priors that are then applied in a Bayesian fashion for further likelihood calculations and iterated until convergence or a maximum of 10 iterations utilizing an Expectation-Maximization schema.

### Validation and sequencing experiments

NumtS identified by computational analysis were validated by polymerase chain reaction (PCR) and Sanger sequencing of amplicon(s) that spanned 50-500bp of gDNA flanking the insert, the breakpoint between the gDNA and the insert, and the insert. Primer sets that hybridize to the gDNA flanking the insert were designed using Primer3 Software (http://www.genome.wi.mit.edu/cgi-bin/primer/primer3_www.cgi) and amplification was done with Platinum Taq (Invitrogen Life Technologies, Gaithersburg, MD), Picomaxx (Agilent Technologies, Palo Alto, CA), or LongAmp (New England Biolabs, Beverly, MA) products in a 20-50ul reaction volume containing 50 ng of template DNA, 1 uM primer, and 1.5mM MgCl_2_ if not supplied in the PCR buffer. Thermocycling was done for 30 cycles at 56-67 °C annealing temperature and 1-15 minute extension time. For inserts less than 3kb, a PCR product of the predicted size was identified in individuals homozygous or heterozygous for the insert by agarose gel electrophoresis and the insert was sequenced in one individual. Amplicons of interest were purified from a PCR reaction for homozygous individuals (Qiaquick PCR purification kit, Qiagen, Valencia, CA) or isolated from the gel for heterozygous individuals (Qiaquick Gel Extraction Kit, Qiagen) and sequenced at the University of Michigan Sequencing Core. For inserts larger than 3kb, a PCR product of the predicted size was identified in individuals heterozygous for the insert by gel electrophoresis. For sequencing, two overlapping PCR products were made using primer sets designed as outlined above with one primer that binds in the gDNA flanking the insert and one primer that binds in the middle of the insert. Amplicons were purified from PCR reactions as outlined above and sequenced by primer walking at the University of Michigan Sequencing Core. Five loci failed initial validation efforts, likely due to a greater than predicted insertion size and uncertainty in the insertion breakpoints. For these loci, we performed a local assembly of the supporting reads using CAP3 (Huang and Madan 1999) and designed additional PCR primers flanking the genome-insertion junction.

### Enrichment analysis

We analyzed the genomic context of the regions flanking the Numt insertion positions for various characteristics, including genes, %GC content, open chromatin regions, repetitive elements, CpG Islands and microsatellites. With the exception of %GC and AT dinucleotide calculations, which were derived from the reference sequence itself using a custom PERL script, all data sets were downloaded from the UCSC Genome Browser (http://genome.ucsc.edu/) in BED format (see Supplemental Table 3 for the specific tables used). We then performed a two-tailed permutation test by resampling one thousand sets of random positions matched to our insertion set to determine whether they were significant enriched or depleted for each feature.

### Phylogenetics and age estimation

An inferred ancestral mitochondria sequence was obtained from ENSEMBL Compara Release 71 based on alignment of six primate species (Flicek et al. 2014). A profile of nucleotide changes was obtained by aligning this sequence to the mtDNA genome from current human reference hg19 using MEGA v5.2.2 with the Muscle algorithm (http://www.megasoftware.net). The age of each NumtS insertion and human-specific reference NumtS (>300bps) was calculated by aligning each sequence to the previously aligned ancestral and modern mtDNA sequence. We tabulated the total number of sites in the aligned region where the ancestral and modern mitochondrial sequences differ and counted how often the NumtS sequence matched the modern human allele. We used the resulting allele matching ratio as an estimate of the point along the human lineage where the insertion occurred.

MEGA v5.2.2 (http://www.megasoftware.net) was used with our larger Numt insertions and previously reported human-specific NumtS present in the reference (Jensen-Seaman et al. 2009; Calabrese et al. 2012; Lang et al. 2012) to determine the evolutionary phylogeny of each sequence. Mitochondrial genomes were obtained for Human (Hg19), Chimpanzee (PanTro4), Gorilla (Gorgor3.1), and Orangutan (PonAbe2) from the UCSC Genome Browser, as well as mitochondrial sequences from Neanderthals (NCBI Accessions: KC879692, FM865411, FM865410, FM865408, FM865407), old European fossil (Homo heidelbergensis) (NCBI Accession: NC_023100), and Denisova (NCBI Accession: FN673705) that were downloaded from the National Center for Biotechnology Information (http://www.ncbi.nlm.nih.gov). The NumtS sequences were aligned to these mtDNA reference sequences using Muscle alignment tool (Edgar 2004) and the trees were built using Maximum-Likelihood method with bootstrap values.

### Haplotyping and Analysis of Heteroplasmy

Single nucleotide polymorphism profiles for each inserted sequence were generated using mtDNAprofiler (Yang et al. 2013). The identified variants were then annotated using Haplogrep (Kloss-Brandstatter et al. 2011) to identify potential haplotypes of origin. These SNP profiles were also used to assess potential heteroplasmy artifacts by direct comparison with reported events (He et al. 2010; Li et al. 2010; Ramos et al. 2013; Bintz et al. 2014; Diroma et al. 2014; Hodgkinson et al. 2014; Jayaprakash et al. 2014). To be considered a match, both the position and the Numt allele must match what was reported, and sample specific information was also indicated where available (Diroma et al. 2014).

## SOFTWARE AND DATA AVAILABILTY

The genomic locations and sequences for the identified NumtS are provided as supplementary data to this manuscript. Sequences of mitochondria insertions and immediate flanking regions have been submitted to GenBank (Accession: KM281512-KM281534). The software package *dinumt* is available for download at https://bitbucket.org/remills/dinumt.

## ACKNOWLEDGMENTS

We would like to thank Xuefang Zhao for her help with figure construction. This project was supported in part through funds from the University of Michigan, the NIH/NHGRI (1R01-HG007068-01A1), and NIH/Common Fund (DP5OD009154).

## AUTHOR CONTRIBUTIONS

G.D. and R.E.M conceived the project idea. G.D. and R.E.M. developed the software, implemented the method and conducted the analyses. S.B.E. and J.M.K. conducted the validation experiments. J.M.K. aided in the age determination and helped with manuscript revision. G.D. and R.E.M wrote the manuscript and prepared figures and tables. R.E.M. supervised the entire project.

